# Predicting the time of entry of nanoparticles in cellular membranes

**DOI:** 10.1101/619221

**Authors:** Changjiang Liu, Paolo Elvati, Sagardip Majumder, Yichun Wang, Allen Liu, Angela Violi

## Abstract

The understanding of the molecular interactions between nanoparticles (NPs) and biological systems is crucial for the systematic advance in many high-impact fields, such as biomedicine and nanotechnology. A key aspect to understand and predict the biological effect of NPs, *e.g*., cytotoxicity, bioavailability, is their interaction with membranes, specifically the mechanisms that regulate passive transport, which controls the permeation of most small molecules. In this paper, we introduce a new streamlined theoretical model that is able to predict the interactions between NPs and biological membranes (average permeation time), by separating the NPs’ characteristics (*i.e*., size, shape, solubility) from the membrane properties (density distribution). This factorization allows the inclusion of data obtained from both experimental and computational sources, as well as rapid estimation of large sets of permutation in new membranes. We validated our approach, by comparing our prediction for the interactions between different carbonaceous NPs and lipid bilayers with both experiments of measuring graphene quantum dot leakage encapsulated in lipid vesicles and time of entry from MD simulations.

## Introduction

The transport of substances across lipid membranes is a biological process of great importance. Molecules that can readily cross cell membranes are frequently needed in biological research and medicine. Permeable molecules that are useful for biological research include for example protein inhibitors, crosslinking molecules, indicators of ion concentrations, and various fluorescent dyes. In medicine, numerous drugs are small molecules acting on intracellular targets, and for most of the routes of administration, cell membrane permeation is required for a drug molecule to reach the internal of the cells. In the case of small organic compounds, it is possible to tune their physicochemical characteristics to facilitate passage through the plasma membrane. Lipophilicity can be increased by avoiding the presence of ionizable groups, and the number of hydrogen-bonded donors can be reduced by N-or O-alkylation. However, this is not the case for most macromolecules, such as proteins or NPs, whose hydrophilicity and large size hampers direct diffusion through the membrane lipid bilayer.^1^

These limitations are especially frustrating in biomedicine. Indeed, recently there has been a blooming of applications in the biomedical arena due to advances in the science of synthesis and characterization of nanoscale engineered particles. Tunable geometric, optical, and surface properties of organic and inorganic nanomaterials enable engineering for a number of applications, such as drug delivery,^2–4^ controlled release,^3,5^ deep tissue imaging and sensing of cellular behavior. ^6–10^ NPs can be delivered by various entry routes, including oral administration,^11,12^ vaccination^13^ and aerosol-based drug delivery,^14–16^ depending on the therapeutic requirement.

However, one of the crucial steps of NP uptake is transport by a physiological cellular membrane. Lipid bilayers are effective barriers to passive diffusion of ions and hydrophilic small molecules, such as carbohydrates, but many molecules can permeate bilayers through passive diffusion at rates that depend on bilayer composition and properties of the permeating solute. The semipermeable nature of the membrane results in an effective selectivity, where small apolar compounds can cross the membrane at appreciable rates. In contrast to trans-membrane ion channels and transporters that are controlled by the cell, passive selectivity is not actively regulated but instead arises from the forces and fluctuations present across the membrane environment. In general, cells do not permit access of polar macromolecules to their cytosol, and phospholipid membranes constitute an effective barrier. Nonetheless, over the years a number of systems have been reported to give cytoplasmic access to biomacromolecules, most notably cell-penetrating peptides^17^ supercharged proteins^18,19^ and bacterial toxins and different types of NPs.^20–22^

Experimental permeability measurements have been performed mainly for many small molecules through various lipid membranes. However, experimental approaches cannot provide adequate information on the mechanism of passive transportation at the molecular level. For macromolecules, such as NPs, NP behavior in biological matrix depends not only on the type of NP, but also on the size, shape, phase, surface charge, chemical composition and agglomeration state.

Over the years, several models have been proposed to describe the process of passive permeation of NP entering the cellular membrane, from the late 19*th* century solubility diffusion model^23,24^ to its recent derivations such as triple-layer model,^25^ MD based approaches^26–28^ and state-transition model.^29^ All these approaches, despite being successful in many applications, often incur large prediction errors for particles of varying size and shape, ^30^ membranes of different compositions^31^ or when membrane phase-changes are involved.^32^

Another group of works instead focused their investigation towards the interaction between lipid heads and NPs, as this is identified as the major factor that controls the permeation barrier.^27,33,34^ This head-group gating effect was theoretically studied in,^30,32^ where the concept of free surface area mediating the permeation was suggested for the first time. Vamparys *et al*. furthermore provided a molecular insight into multiple definitions of the free surface area as well as the area distribution.^35^ While promising, the free surface area model, is still hard to apply as it has several critical shortcomings: not only does this model exclude the shape, but also the lifetime distribution of the free surface areas. As we will show later, these characteristics are essential for building a generic permeation model.

In this paper, we present a new theoretical model that predicts the average time of permeation of NPs through the hydrophilic region of a lipid membrane, using a combination of molecular dynamics (MD) and ad hoc analysis. The most important contribution of this new model is the identification of four parameters to describe the process and the factorization of the contributions that are only dependent on the characteristics of NPs (e.g., size, shape) from the ones that depend on the properties of the lipid membrane (e.g., free surface area) and the one that depend on both (solubility). This separation has two main benefits: it allows to use data obtained from both experimental (e.g., partition coefficients, NPs shape measures) and computational sources (membrane surface dynamics) as they become available, and it allows a rapid estimation of large sets of new systems by using existing data. The new model is validated using both experimental data on vesicles and NPs and all-atom MD simulations, obtaining in all cases agreement well within measurement error. Given the high level of interest across multiple areas of study in modulating intracellular targets, and the need to understand and improve the applications of NPs and to assess their effect on human health (i.e., cytotoxicity, bioavailability), this work contributes to the understanding and prediction of time of entry of NPs into lipid membranes.

## Results and discussion

### The Permeation Process

To identify trends in membrane permeation that can be used for our model, we performed several independent MD simulations of different NPs in a phosphatidylcholine (POPC)/cholesterol bilayer (10:1 concentration ratio). The ratio represents the typical composition of a mammalian cellular membrane. ^36^

As for NPs, we focus our study on carbonaceous NPs since they have been widely utilized in many health-related fields like drug delivery,^2–4^ cancer therapy,^3,5^ wound disinfection,^37^ biolabeling^6–8^ and biosensing,^9,10^ due to their low cytotoxicity, high photoluminescence and easy surface functionalization. Specifically, a buckminsterfullerene, a curved OH-terminated graphene quantum dot (GQD) and GQD functionalized with two cysteine groups (cys-GQD) were used.^38^ This selection covers NPs of similar size, but different shape and hydrophilicity.

Analysis of MD trajectories show that the process of permeation can be divided in four stages, summarized in Fig. 1A,B: **I.** The NP reaches the membrane surface via diffusion. The NP diffuses over the water-membrane interface. **III.** The NP enters the hydrophilic region of the lipid bilayer. **IV.** The NP permeates the hydrophobic region of the lipid bilayer. These stages are not strictly sequential. For example, from Stage II, the NP can go to either Stage I or III, or, depending on the NP’s properties, Stage II or III may be a long-lived state that does not further evolve in a given time frame.

**Figure 1:**
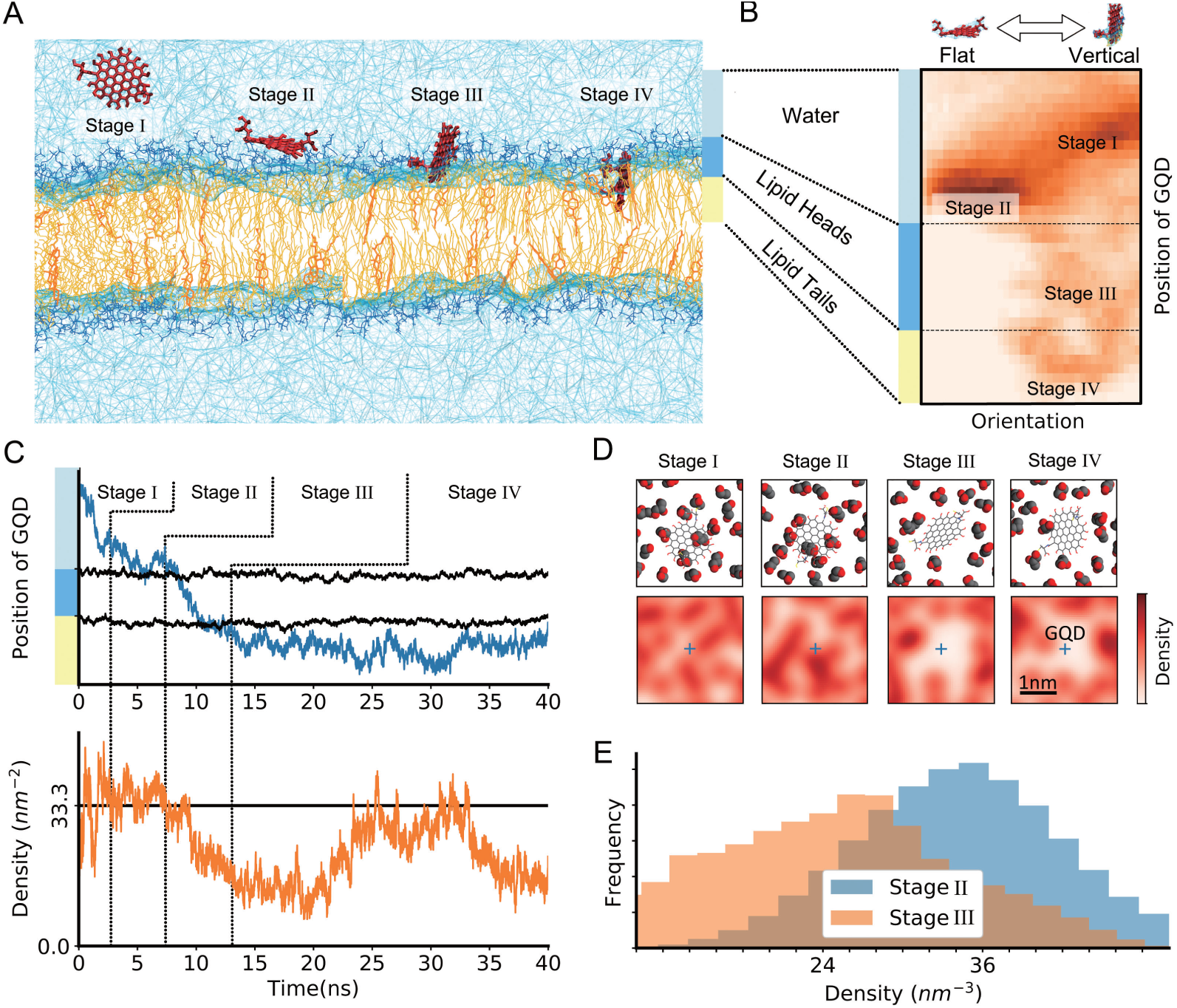
Permeation of a 2-nm cys-GQD in a lipid bilayer. The process can be separated in four phases (A), in which position and orientation of the permeant strongly correlate (B). Lipid head-groups positions and surface lipid density (D) show the LDA in the lipid-head region formed around the GQD during stage III. This effect can be also seen when comparing the time evolution of the cys-GQD distance from the membrane central plane (C, upper panel) with the average atomic density of the membrane surface around the GQD (C, lower panel), as well as the average distribution of the density around GQD during stage II and stage III (E).

The first transition (from Stage I to Stage II) is controlled in most conditions by the diffusion of the NP in the environment close to the membrane surface, due to the presence of an unstirred layer.^39^ As this environment can be strongly affected by the species (*e.g*., proteins and saccharides) that surround many biological membranes, and it is largely independent of the bilayer composition, we decided to restrict the focus of our model to the other transitions.

When analyzing the progress from Stage II to Stage III, we consistently observed a drop in the membrane surface density in proximity of the particle (Fig. 1C-E). At the same time, for non-spherical NPs (*e.g*., cys-GQD), the transition is characterized by a change in the orientation of the NP, namely from parallel to perpendicular to the membrane plane (Fig. 1B), in agreement with previous works.^38,40,41^ In order to determine whether the decrease in local density of membrane surface is caused by the presence of the NP or is independent of it, we computed the membrane density distribution and dynamics both in absence and presence (close proximity) of NPs.

The result (Fig. 2) shows that neither the density distribution nor the autocorrelation is significantly affected by the presence of the NPs selected in this work, as the difference between the curves (purple) is always smaller than the standard error (dashed, red). This result suggests that the local change in density is due to the internal dynamics of the lipid bilayer, *e.g*., thermal fluctuations. It is important to note that we do not exclude that different NPs can affect the local membrane density, most likely if strong Coulombic interactions are present. This is the case in compounds like graphene oxide nanosheets and a wide variety of cationic NPs, which are known to induce pore formation, membrane thinning and membrane erosion. ^40,42–44^ However, as we will show below, the effect on permeation due to induced membrane changes can be modeled as a subset of the more complex case of fluctuations driven density changes that we observed.

**Figure 2:**
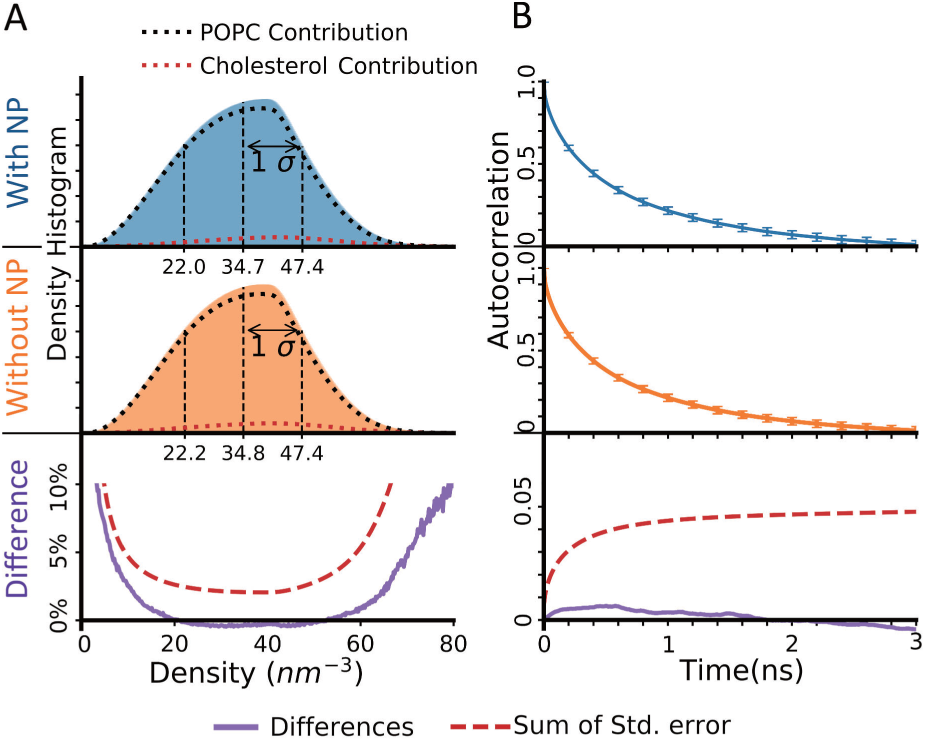
Effect of the presence of NPs (three different NPs, see text for details) on a POPC/cholesterol (ratio 10:1) bilayer density. Error bars (95 % confidence interval) shown only for selected points for clarity.

### The Model

Based on the insights gained from MD simulations illustrated above, we developed a simplified model for NP permeation in biological membranes. The model describes the permeation as a gated entry model, in which the single process is regulated by the chance of the NP to encounter low-density areas (LDAs) on the membrane surface and by the solubility of the NP in the hydrophobic phase. Once the NP reaches the membrane surface, the process can be summarized as follows (Fig. 3):

**Figure 3:**
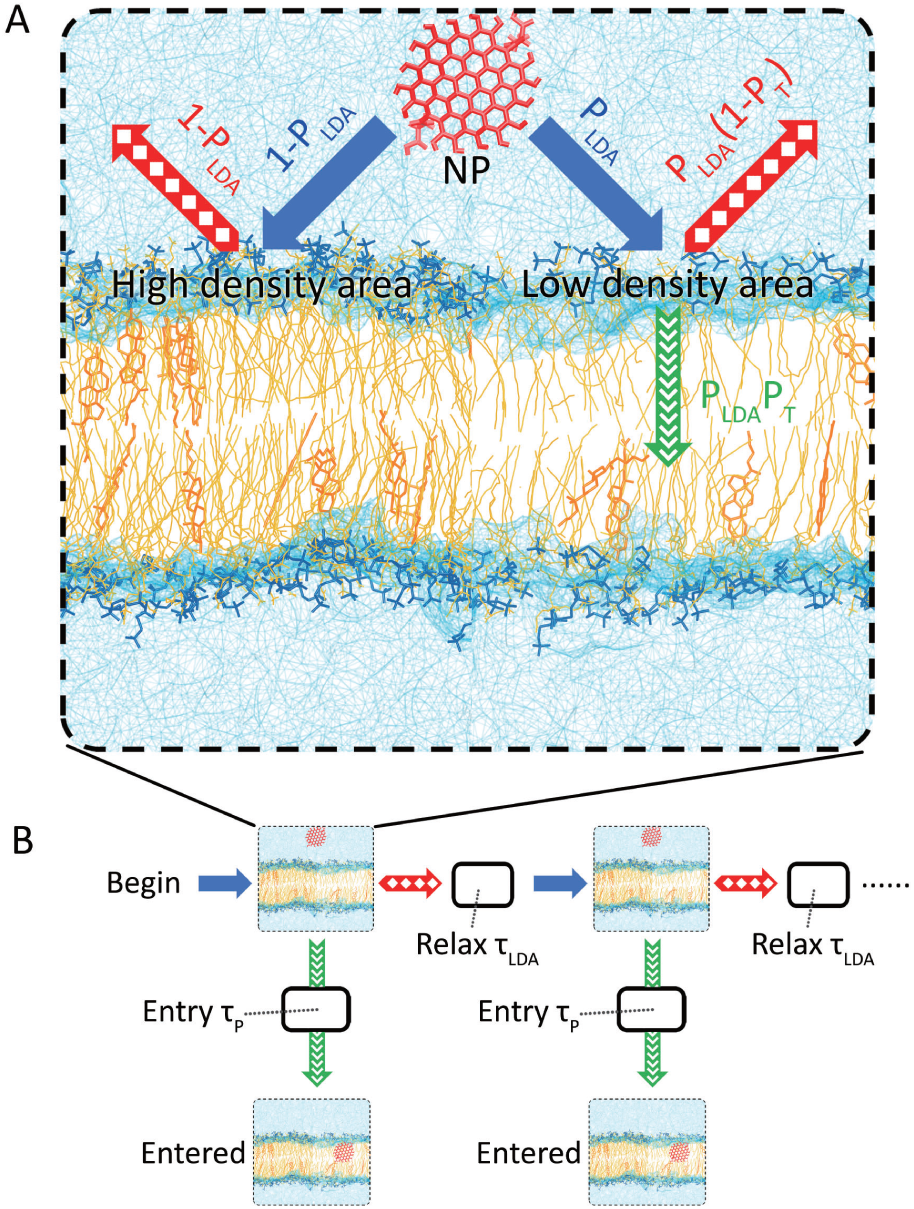
Schematic representation of the proposed permeation model. (A) The probability of entry controlled by *P*_*LDA*_ and *P*_*T*_. (B) The diagram of time evolution. See text for the detailed description.

1. The NP that is in proximity of the membrane surface may encounter a local LDA of lipids with the right shape to allow the NP to permeate, with a probability *P*_*LDA*_. *P*_*LDA*_ is generally dependent only on the membrane properties (*e.g*., composition, surface tension). For NPs that perturb the local density this probability should be modified.

2. If the LDA is large enough to allow NP permeation, the process is now controlled by its affinity with the lipid tails, *P*_*T*_.

3. If permeation does not occur, the NP can leave the membrane surface or stay in proximity of the membrane long enough for a new independent “permeation attempt” (two previous steps) to occur. To approximate the time elapsed between two independent permeation attempts, here we use the LDA decorrelation time, *τ*_*LDA*_, during which the thermal fluctuations generate a new decorrelated distribution of LDAs on the membrane. Moreover, if we assume a condition of steady state, the probability of a NP diffusing away from the membrane is the same of a different NP reaching the membrane.

Based on this description, the average time of entry *t*_*entry*_ can be estimated as:

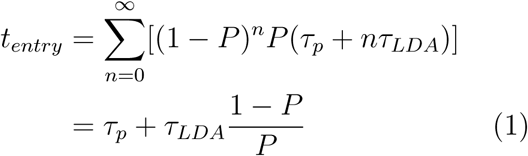

where *τ*_*p*_ is the average time for the NP to diffuse from the lipid head region to the lipid tail region and *P* is the probability defined as:

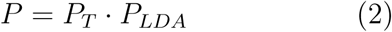

Below we discuss in details all the four parameters reported in Eq. 1 and Eq. 2.

#### Thermodynamic probability (*P*_*T*_)

The affinity of particle with the lipid tail *P*_*T*_ describes the probability of the NP to move from the hydrophilic (headgroups region) to the hydrophobic phase. This factor is therefore related to the solubility of the NP in the membrane and can be estimated using the partition coefficient *K*_*D*_ of the NP in water/lipid as:

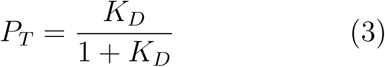

This term agrees with the observed strong correlation between the permeability of NP in different biological membranes and the NP water/membrane partition coefficient.^27,45,46^

*K*_*D*_ can be obtained from both computational and experimental techniques or approximating with simpler system, *e.g*., the water/octanol partition coefficient. Multiple theoretical models have been proposed also to estimate the water/octanol partition coefficient. For examples, Garrido *et al*.^47^ used MD simulations to determine the temperature dependent partition coefficients of different *n*-alkanes obtaining a good agreement with experimental values. Theoretical methods that do not require simulations^48,49^ can also be used to predict the partition coefficient using only the atomistic and structural information of a particle.

#### LDA probability (*P*_*LDA*_)

*P*_*LDA*_ is the probability of finding an LDA of a given size and it is affected by both the properties of NP and membrane. However, for NPs that do not markedly alter the membrane surface dynamics, the contributions of membrane and NP can be separated and the *P*_*LDA*_ can be computed from the size and shape distributions of the LDA in the membrane and the size and shape of the NP, as described below.

The LDA distributions can be computed in different ways, and below we propose an approach that can be extended potentially to experimental techniques, by converting atomistic three-dimensional information to two-dimensional images, which in turn are used to compute the LDA properties.

Next, we illustrate all the steps of the procedure by using the data from a MD simulation of a POPC:cholesterol (10:1) bilayer at physiological ionic concentration (0.15 M NaCl).

For each sample of our simulation, we located the center of mass (COM) of each lipid by considering only the atoms at the water interface (Fig. 4A). The membrane surface is then segmented using the COM positions to build a Voronoi diagram (Fig. 4B). This partition makes it possible to determine the average area “occupied” by each lipid type under the specific environmental conditions and membrane composition. To obtain a continuous density map, we summed the two-dimensional Gaussian function centered at each lipid COM with a radius dependent on the average membrane area for that type of molecule, *e.g*., cholesterol, POPC (Fig. 4C). With these steps, we have transformed the three-dimensional information from atomistic simulations to a continuous two-dimensional density map of the water/membrane interface, similar to the one that can be obtained from different imaging techniques (*e.g*., atomic force microscopy, electron microscopy). Therefore, the following steps can be in general applied to any two-dimensional density map independently from its source.

**Figure 4:**
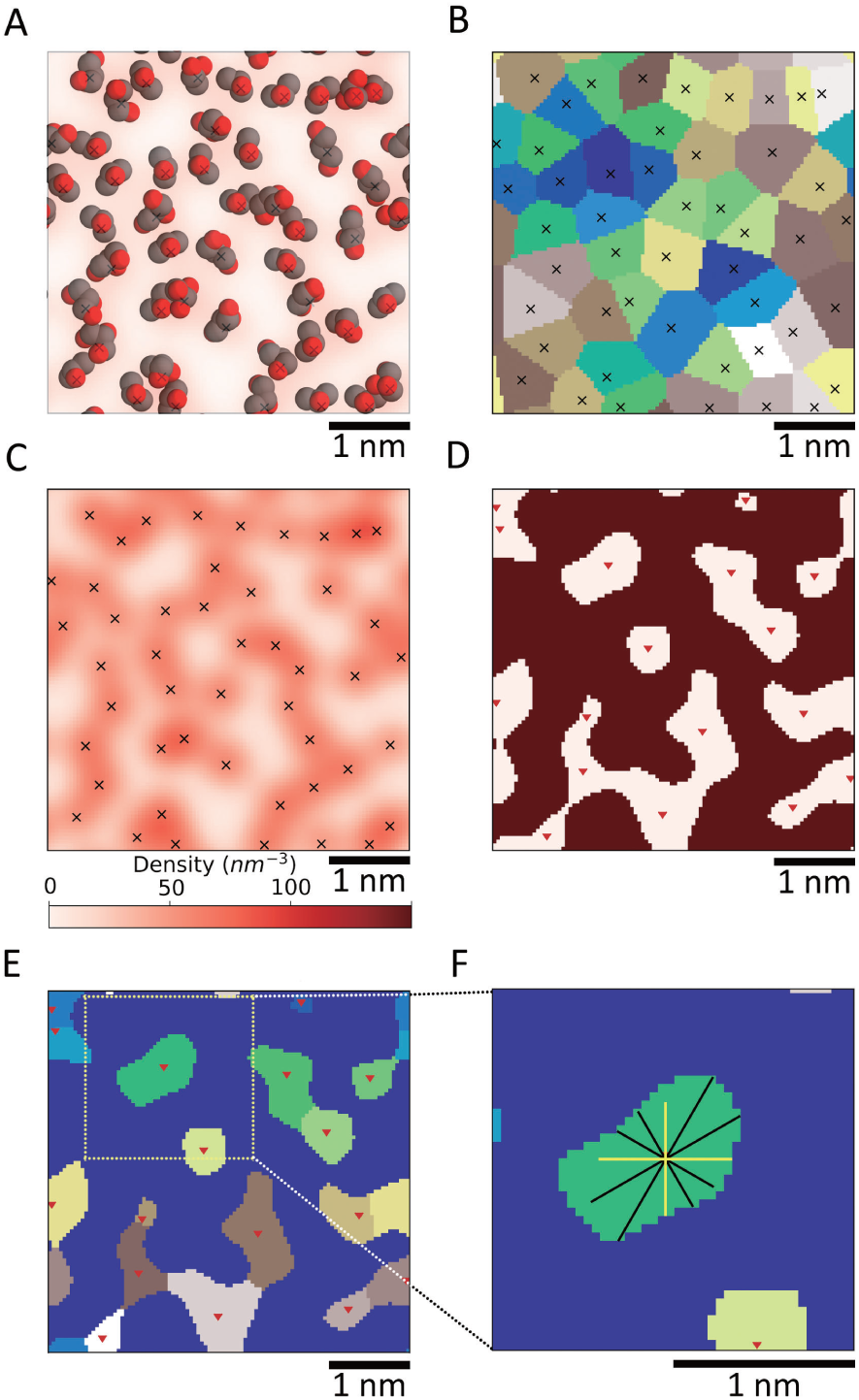
Schematic representation of the steps used in the analysis of the lipids’ density for each sample of a POPC:cholesterol (ratio of 10:1) bilayer (top-down view). See methodology for additional details. (A) Calculation of lipids’ COM; atoms at the lipid/water interface are selected to locate the center of mass for each lipid. (B) Voronoi segmentation; crosses show lipids’ COM. (C) Continuous density map obtained by applying a Gaussian function centered at each lipid COM. (D) Regions with a density above (light red) and below (dark red) the average water density (33 nm^3^ excluding hydrogen atoms). Down triangles indicate the local density minima. (E) Segmentation of joint LDAs by watershed by flooding on the density contour. (F) Dimensions probing of a single LDA, by sampling on perpendicular directions (*e.g*., yellow lines) passing through the LDA minimum (only 3 pairs are shown for clarity).

The next step is to extract the statistics of LDAs’ morphology from the two-dimensional density map. We first assign the center of all the LDAs to the local minima in the two-dimensional density (Fig. 4C); the LDAs are then extended from these centers to the contour line corresponding to the density of water, *i.e*., 33 nm^3^ (Fig. 4D). Finally, joint LDAs are separated by a flooding watershed algorithm^50^ (Fig. 4E).

As shown in Fig. 4E, the shapes of LDAs produced from the segmentation method, described above, are often irregular. To describe these shapes, we draw perpendicular lines starting from the local minimum at 18 different angles (Fig. 4F). The lengths of these perpendicular lines are then used as descriptors of the shape and size of an LDA. Specifically, these dimensions can also be used as a filter for the size of the permeating particle.

The average filter over all the angles is computed for each LDA and then weighted by the LDA area. The result, normalized by the total membrane area, gives the size dependent permeation probability for the given two-dimensional density map. Finally, the average over all the samples (*i.e*., the MD trajectory) gives *P*_*LDA*_ as a function of the particle dimensions. One example shown in Fig. 5.

**Figure 5:**
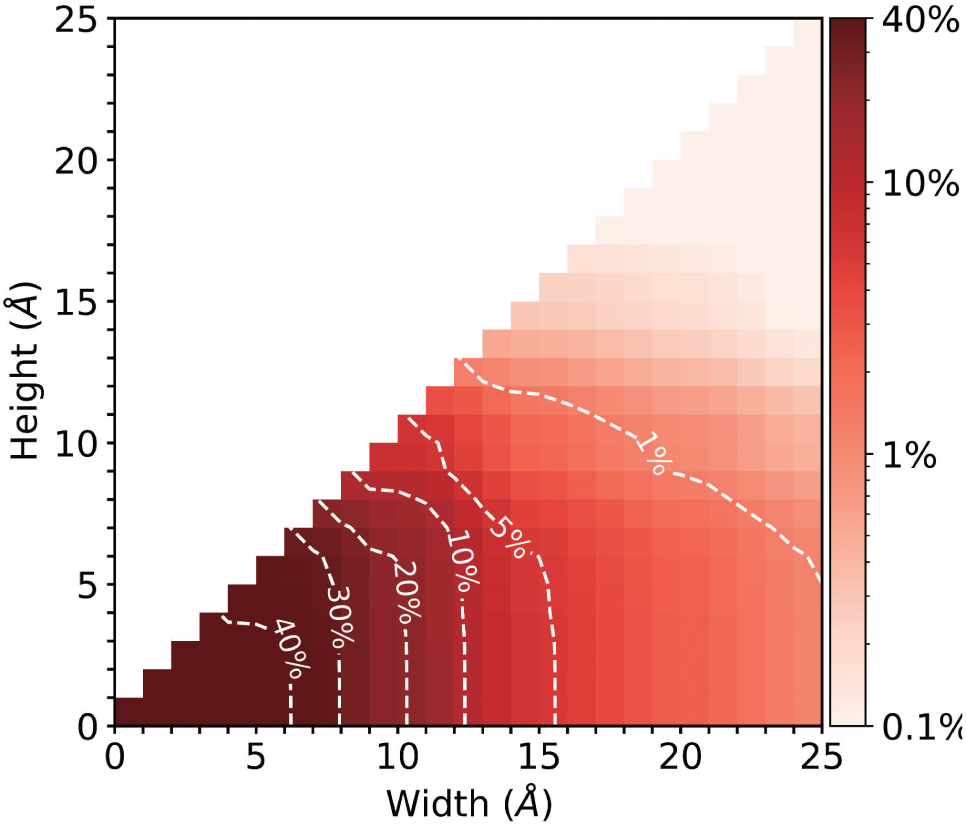
Average probability of finding an LDA larger than a given rectangular cross-section, shown as cumulative histogram in a POPC/cholesterol (10:1) membrane.

To account for the different possible orientations of a particle, we considered a cross-section along the 6 directions defined by the particle’s axes of inertia.

To eliminate artifacts due to barbed or oddly-shaped objects, the cross-section was taken 0.28 nm from the particle surface, approximately the thickness of one water layer, and then approximated by its circumscribing rectangle. Finally, the sizes of the rectangles are used to compute six values of *P*_*LDA*_ that as an average represent the overall particle *P*_*LDA*_ but can also be used to estimate the preferential orientation during permeation.

#### LDA decorrelation time (*τ*_*LDA*_)

For the model to include also information on the dynamics of the membranes, knowledge on the time evolution of the LDAs need to be included. *τ*_*LDA*_ is the average time that the membrane surface density takes to decorrelate, which is controlled only by the membrane thermal and pressure fluctuations, under the non-interacting case discussed above. We estimated the *τ*_*LDA*_ using the thresholded two-dimensional density map (*e.g*., Fig. 4D), and measured the shortest time of part of the LDA to transition to a non-LDA. As a measure of the decorrelation, we approximated *τ*_*LDA*_ as the characteristic time needed for the 95% conversion from LDA to non-LDA.

Fig. 6 shows the distribution of the lifetimes and *τ*_*LDA*_ for three different surface density thresholds. While there are differences in the curves depending on the threshold value, once a 93-97% conversion interval is considered, taken into account the different sources of error, the results are substantially similar with values close to 2 ns.

**Figure 6:**
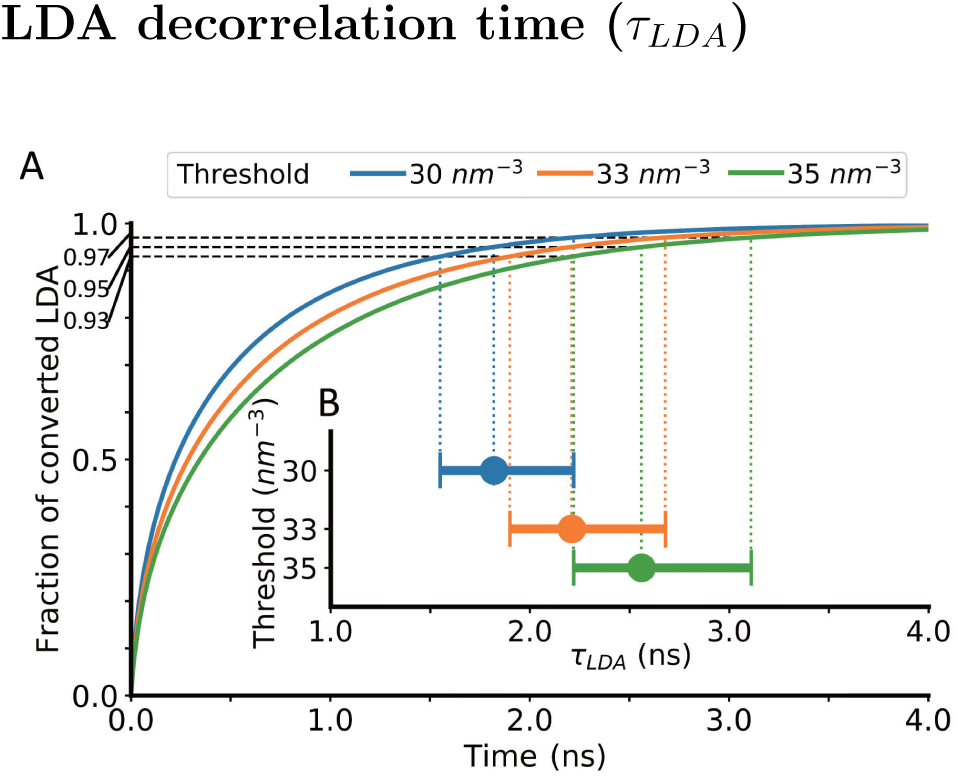
Estimated lifetime of LDA, based on the (A) fraction of LDAs that converts to non-LDA over time. (B) *τ*_*LDA*_ computed for different thresholds, for 95% LDA lifetime (bar indicate the 93-97% interval).

In this work *τ*_*LDA*_ is extracted from surface density maps, but it can be derived also from theoretical models or experimental measurements of the fluctuations of the membrane surface.^51,52^ An intuitive comparison of the order of magnitude of the lifetime of LDA to membrane lipid dynamics are lipid wobble (ns), axial rotation (ns) and lateral diffusion (ns to ms).

#### Time of permeation (*τ*_*P*_)

*τ*_*P*_ is the average time that takes the NP to permeate from the lipid heads to the lipid tails when no barrier is encountered. This time provides a small contribution to the overall time of entry as compared to *τ*_*LDA*_, as the permeation is limited by the head groups. Numerically *τ*_*P*_ can be approximated based on the diffusion velocity of the NP, estimated from the size and mass of the NP,^53^ but in this work we used the difference in time between the formation of an LDA and the time of entry from the MD trajectories. This approach leads to values approximately between 1 ns for C60 to 3 ns for cys-GQD (refer to Fig. S3 in the supporting information).

### Comparison between LDAs and packing defects

The LDA is not the only way to evaluate the tightness of lipid packing at the hydrophilic/hydrophobic interface of lipid bilayers. There are also other metrics, such as accessibility surface area^54^ (which uses a sphere to probes the void on membrane surface) and packing defects^35^ (which calculates the void by projecting the membrane surface to a grid). In the work done by Vamparys *et al*.,^35^ these metrics are compared accordingly to their distributions of the sizes of defect areas. The results showed that, for all these metrics, 1) the sizes of defect areas increase when packing defects are introduced into the lipid bilayer, and 2) the distributions of the sizes of defect areas are roughly exponential.

Due to the conceptual similarity between LDA and the defect area, here we compare them, and following the approach as introduced by Vamparys *et al*.,^35^ we computed the chemical and geometrical defects for three different membranes varying their POPC/cholesterol ratios. Analyses are performed on 2500 snapshots from 50 ns simulation for each membrane as shown in Fig. 7. For all lipid compositions, we observe the exponential decay in size for all the distributions, above a minimal size (0.05 nm^2^ for defects, 0.5 nm^2^ for LDA) similarly to the results presented in the original work.^35^ This decay increases with the amount of cholesterol in the membrane, implying that the cholesterol facilitates the lipid packing in the bilayer, resulting in less packing defects on the membrane surface, in agreement with the previous works.^55–58^ These similarities indicate that the metric of LDAs captures similar membrane surface properties as the metrics of packing defects do.

**Figure 7:**
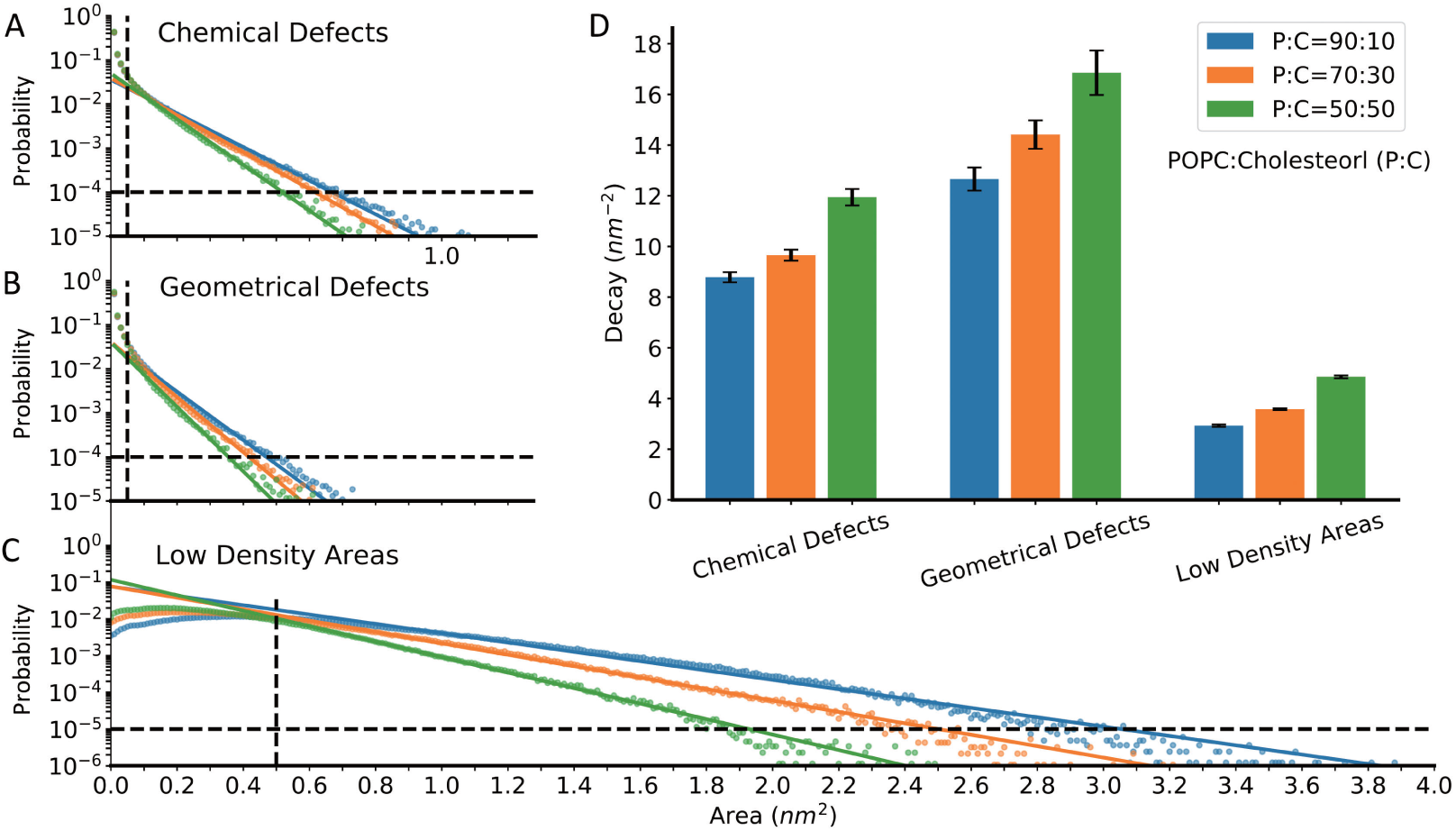
Size distribution of packing defects and LDAs in model membranes. The distribution is the probability of finding a defect area or LDA of defined size. Panel (A, B) are for the chemical defects and geometrical defects introduced by Vamparys *et al*.^35^ and panel (C) for LDAs introduced in this work. For each metric, we analyze three types of membrane varying in their composition (ratio of POPC:cholesterol). Colors blue represent the POPC:cholesterol=90:10, orange for 70:30, and green for 50:50. Dots represent each sample and solid lines show the exponential function fit. Following the same approach as the original work by Vamparys *et al*.,^35^ to fit the exponential function for the distribution of the two types of packing defects, we selected only defects larger than 0.05 nm^2^ and with probability greater than 10^*-*4^. For LDAs, this range of selection is for area larger than 0.5 nm^2^ and with probability greater than 10^*-*5^. The black dashed lines in panel (A, B, C) indicate such ranges of selection. Panel (D) compares the exponential decay constants for all the membranes and all the metrics.

However, the data in Fig. 7 highlights also some of the differences between these definitions. The most important one is the shape of the curve approaching zero: while the packing defects monotonically decrease with increase defect size, the LDA distribution shows a maximum around approximately 0.5 nm^2^. The reason for this difference lays in the physical meaning of the LDA, namely the free energy cost associated with the membrane density fluctuations. On one hand, any deviation from the average density is not entropically favored. On the other hand, the fluctuations are enthalpically favored for small changes, as the average lipid-lipid distance becomes closer to the one associated with the potential energy minimum, but it then becomes also unfavorable once the repulsive part of the lipid-lipid interactions become prominent. Finally, the typical size of the LDA is about three times larger than the packing defects. This difference is intrinsic in their different definitions, as the packing defects are measuring the voids between atoms, which is a much stricter requirement than the LDAs defined by a threshold of density.

### Validation

To test the performance of the new model, we carried out two comparisons: the first one against statistical observations of a set of all-atom MD simulations, and then versus experiments. As currently only *P*_*T*_ is estimated from experimental values, the first comparison establishes the ability of the model to capture the MD process independently of the approximations and simplifications introduced above.

To obtain the time of entry of three different NPs (Fig. S1), we ran 12 to 16 simulations by placing the NP in close proximity of a POPC/cholesterol bilayer (10:1 ratio) while immersed in a NaCl solution at physiological conditions (0.15 M). The times of entry were collected and fit with an exponential function from which the characteristic time was determined as the average time of entry. The quality of the fit can be evaluated by the standard error of the time of entry (via its Fisher information). The average time of entry and the standard error are listed in Table 1, along with other parameters used in the model.

**Table 1:**
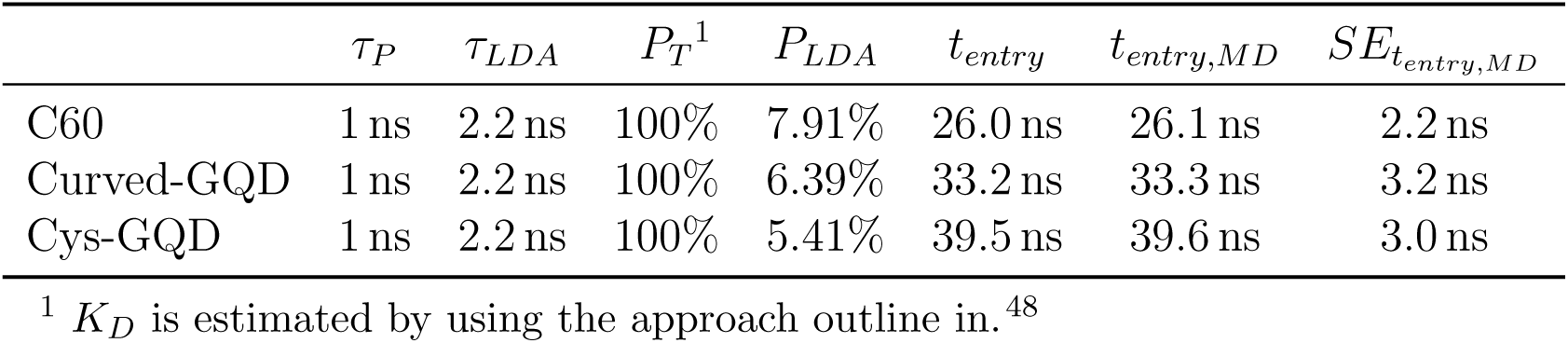
Parameters related to the model’s prediction for the three CNPs.

The results are reported in Fig. 8, which shows that the agreement between the MD and the current model is within 10%. It is worth noting that all the 3 predictions seem to suffer from a negative bias error of about 0.1 ns. Since this term is consistent for all NPs, we attribute that to the factors that depend solely on the membrane, *i.e., τ*_*LDA*_.

**Figure 8:**
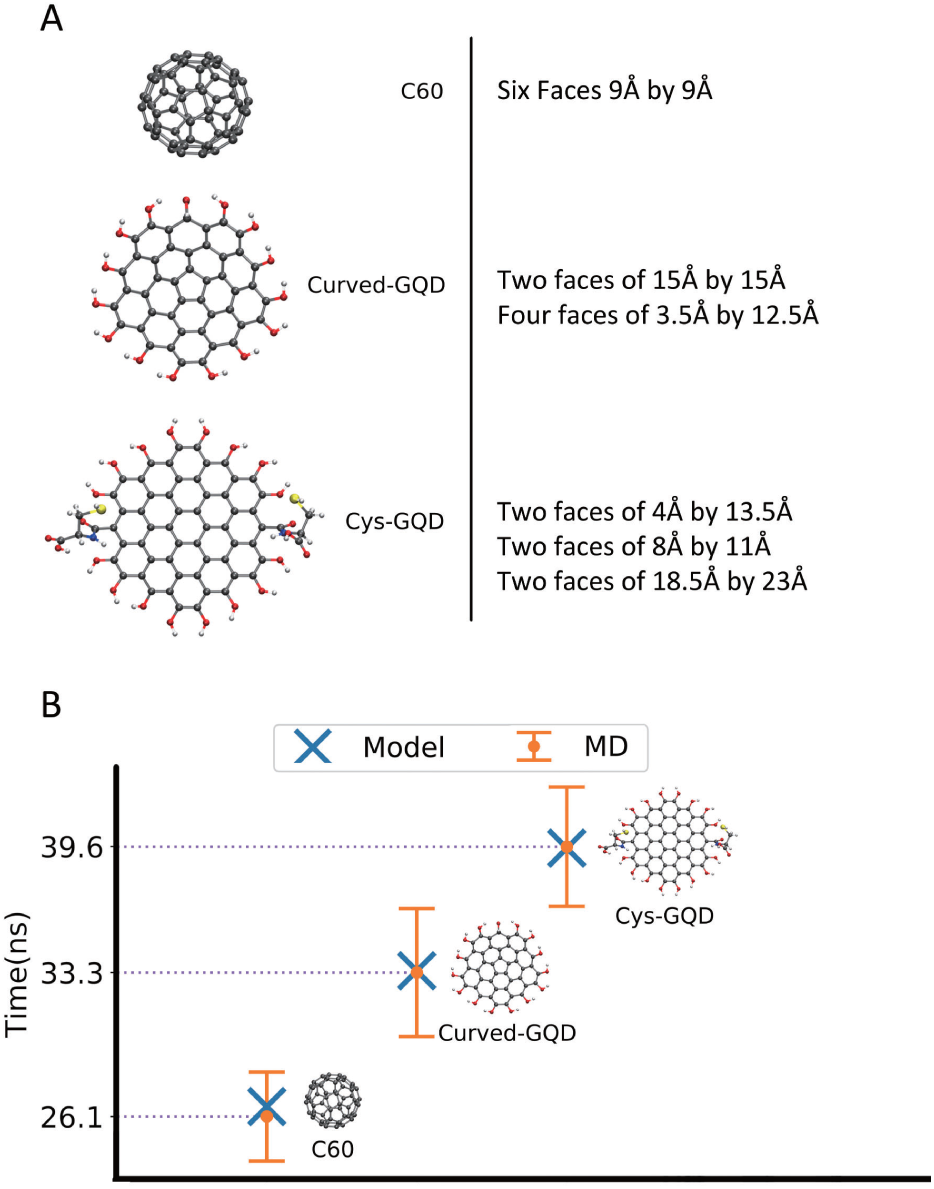
Comparison of the average time of permeation predicted from MD simulations and the current model.

Given the overall excellent agreement between the MD and the model, we tested the latter by comparing it with the experimental measurements of a photoluminescent NP crossing the membrane of lipid vesicles of different compositions. Specifically, we leverage the linearity of the photoluminescence intensity of a GQD solution with its concentration, by monitoring the decrease in luminescence of a GQD solution encapsulated in different vesicles, as a measurement of GQD leakage. The lipid vesicles encapsulating with GQD (0.1 mM), with three different POPC/cholesterol ratios (*i.e*., 50:50, 70:30 and 90:10), and GQD ranging from 2 nm to 8 nm in diameter, were prepared by double emulsion templated vesicles. ^59^ Next, the lipid vesicles were moved to a GQD-free solution and the GQD leakage was measured by monitoring the decrease of the photoluminescence of a single lipid vesicle over 1-hour period (Fig. 9A) from the time the vesicle was moved to a GQD-free solution. The results, presented in Fig. 9A,B are the average over 10 independent vesicles.

**Figure 9:**
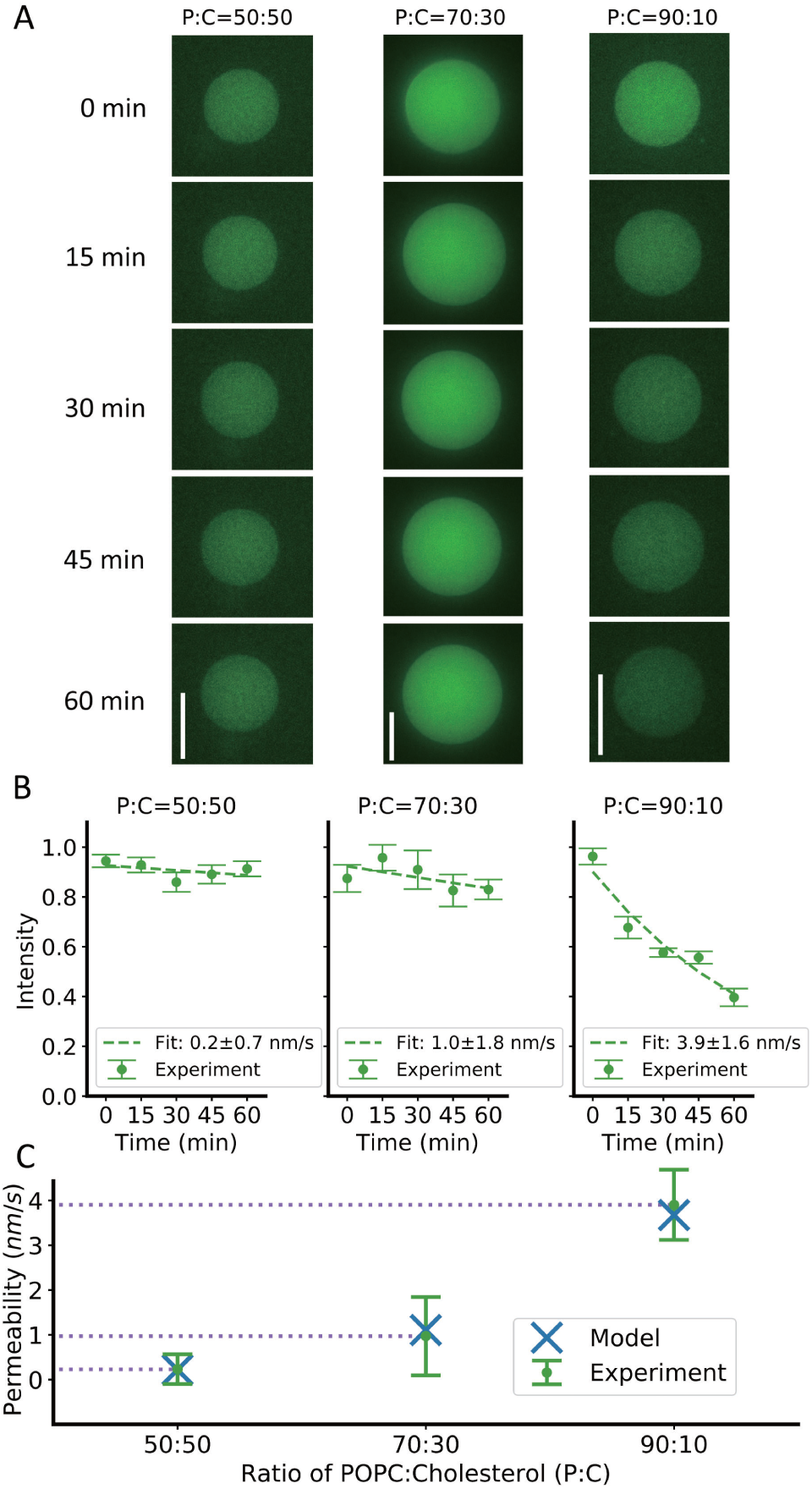
Measured GQD leakage from different lipid vesicles. (A) Experimental images of photoluminescence change over one-hour period. Images were taken every 15 minutes. White scale bar for 50 µm. (B) The photoluminescence intensity over one-hour period for GQD encapsulated vesicles with different lipid compositions are indicated. (C) Comparison of the model’s predictions to the permeability measured from experiment. (Error bars correspond to one standard deviation.)

The kinetic of the GQDs’ leakage in a spherical vesicle can be modeled by a first-order dynamics as

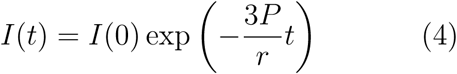

where *I*(*t*) is the intensity of the fluorescence over time, *P* is the permeability of the membrane, *r* is the radius of the lipid vesicle. We modeled the leakage of GQD as a two-step process, where the GQD first enter the vesicle bilayer and then either diffuse back inside or outside the vesicle. If *I*(*t*) is proportional to the GQD concentration inside the vesicle and the bilayer, this model predicts a decrease in intensity dependent on GQD concentration in the membrane [*GQD*]_*M*_:

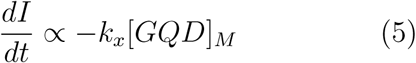

where *k*_*x*_ is the kinetic rate describing the GQD leaving the bilayer. Based on the values of the partition function as well as the difference in timescale between the leakage process and the entry process (seconds or minutes *vs* nanoseconds), we assume that a steady state condition is rapidly reached. Therefore, by comparing Eq. 5 with Eq. 4, we find that *P ∝ rk*_*x*_*/*3 = *c/t*_*entry*_, where *c* is a constant that depends on the membrane thickness, the partition coefficient and the fraction of particles that go outside the membrane when leaving the membrane. As it is shown in in panel c of Fig. 9, there is a very close match between the permeability obtained with our model and the one computed by fitting the experimental data (dash line in Fig. 9), further validating the quality of our approach.

## Conclusion

In this work, we presented and validated a kinetic model for the permeation of NP in cellular membranes that captures the atomistic detail of the process while retaining the simplicity typical of macroscopic models. The model was built on the observation, based on all-atom simulations, that the permeation process is controlled by the formation of LDAs on the membrane surface. The dynamics of these LDAs is generally independent of the presence of NPs, as it was observed for the carbonaceous NP selected for this study, but the model can be straight-forwardly modified for particles that have long range interactions with the membrane, *e.g*., charged NP.

Based on these findings, we factorized the process of permeation into the independent effects of membrane thermal fluctuation and the particle’s characteristics (size, shape and solubility in membrane). A combination of these two factors provides a physical model to predict the time elapsed from the NPs in contact with membrane to NP fully emerged in the lipid tails. Under the assumption of local equilibrium, this time also serves as the time constant for the model of permeation with barrier using the first-order dynamic.

It is the first time that the surface dynamics of the membrane is discussed from a MD perspective. As a physics-based model, this model can be efficiently applied to a wide range of NPs and membranes. The model was tested on CNPs of different structure and the same CNP on different lipid vesicles. Predictions from the model matched quantitatively to the experimental results and simulations, indicating the success of the underlying physical model.

The parameters in the model may be validated through different experimental and theoretical approaches. For example, membrane with increased tension can induce more LDAs and may be used to study the trend of the increase in membrane permeability. Atomic-force microscopy may be used to directly probe the surface atom density distribution. Also, some theoretical model may be suggested to describe the thermodynamics underlying the formation of LDAs.

As the physical model and factorization provided insight into the mechanism of NP/membrane interactions, it also opens a new direction for further research in this field. For examples, understanding the selectivity based on the membrane composition provides information for designing NPs to target specific sites. The efficient prediction made from this model also enables high-throughput drug screening for bioavailability assay.

Some improvement in the future can be made to the model. As our approximation of rectangle boxes for the cross-sectional areas over six characteristic directions was a preliminary one, a full integral over the 3D angles may better utilize the information provided by the structures of NPs. In the meantime, interactions between the membrane and NPs may influence the distribution of the orientation of NPs toward membrane plane, which if taken into consideration, may improve the prediction as well. In addition to mammalian cell membrane, this model may also help learning insights for systems such as bacterial lipopolysaccharide membranes and polymer chains.

## Methodology

### MD Simulations

We used NAMD^60^ version 2.13 with CHARMM Force Field^61^ version 36 for lipids, CHARMM General Force Field^62^ for simulating the CNPs and TIP3P^63^ for water molecules. CGenFF^64^ software was used to assist in the production of the needed topology files. The initial placement of the membranes 120 POPC and 12 cholesterol molecules (60 and 6 on each leaflet, respectively) and 8280 water molecules was performed with the CHARMM-GUI membrane builder.^65^ All MD simulations were performed with a timestep of 2 fs while keeping all the C– H and O – H bonds rigid via the SHAKE algorithm.^66^ Long range electrostatic interactions were modeled with the particle mesh Ewald method^67^ using a 0.1 nm grid spacing, a tolerance of 1E-6 and cubic interpolation, and cubic periodic boundary conditions were applied. Temperature was kept constant at 310 K using a Langevin thermostat^68^ with a time constant of 1 ps, while pressure was kept at 101.325 kPa using Langevin piston method^69,70^ with period of 50 fs and 25 fs decay. To account for the intrinsic anisotropy of the system, the production simulations were performed in the NPsT ensemble, where the x and y dimensions of the periodic system (coplanar with the membrane) are allowed to vary independently from the z dimension. Non-bonded short-range interactions where smoothly switched to 0 between 1 and 1.2 nm with a X-PLOR switch function.

The POPC:cholesterol = 10:1 membrane was equilibrated for a total of 80 ns, while the other three membranes (POPC:cholesterol = 90:10, 70:30 and 50:50) were equilibrated for 130 ns simulations. The membranes were assumed to be at equilibrium once the time average of the area per lipid in a NPsT simulations was varying less than 10%, as shown in Fig. S2. The surface density dynamics for each of these membranes was analyzed on the final 50 ns of the production simulations.

The permeation dynamics were studied by placing the different molecule (2 nm) away from the membrane’s head groups. To stop the molecules from diffusing too far away from the membrane, we imposed a harmonic potential that pushes the particles towards the membrane when its distance from the membrane interface exceeded 2 nm. We run 16 simulations for the cys-GQD, 12 for curved-GQD, and 12 for C60, in each case for either 40 ns or until the molecule was embedded in the hydrophobic region of the membrane for at least 1 ns.

VMD^71^ and MDAnalysis^72,73^ were used for data analyses and visualization.

### Average Time of Entry

To estimate the average time of entry for each CNP, we plotted the evolution of their distance from the membrane center (see Fig. S3) and collected the time of entry. The green (or yellow for non-entry cases) curve tracks the COM of the CNP during the simulation, and the black curve is the COM of the lipid head-groups (including the glycerol group and phosphorous group for POPC, and the 3,4,5-carbons and the oxygen for cholesterol). The time of entry is measured as the time passed from the beginning of the simulation to the intersection of two curves.

Following the assumption that the entrance is a random event, we fitted the collected times with an exponential distribution:

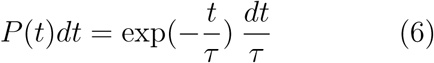

where *τ* denote the exponential parameter that has the same unit as *t*.

In each one of the *N* independent simulations, the NP can either enter the lipid tails region during the simulation (*t*_*sim*_ =40 ns) or not. The likelihood *L* for *n* simulations to satisfy *t*_*i*_*<* 40 ns is given by:

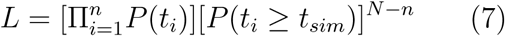

From Eq. 6 and 7, we can find the *τ* that maximize the likelihood:

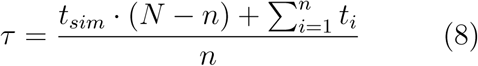

and the expectation and standard error (from Fisher information) of the time are *τ* and 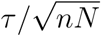, respectively.

### Definition of LDAs

The steps used to define the LDAs and measure their size/shape are presented in the text, but minor additional details are described below.

1. The center for each lipid (Fig. 4A) is calculated using the center of masses of the heavy atoms near the water/lipid interface (glycerol group for POPC, and top 4 heavy atoms for cholesterol).
2. Based on the center of each lipid, the Voronoi map (Fig. 4B) is created to calculate the average area for each type of lipid (POPC: 0.603 nm^2^, cholesterol: 0.452 nm^2^).
3. The continuous density map (Fig. 4C) is created by applying a Gaussian kernel centered on each lipid COM. The height of each kernel is equal to the number of heavy atoms selected for the mapping, while the standard deviation of each kernel depends on the lipid type and is related to its average area by

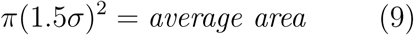

Final density of atoms per area is compared with the density of water of atoms per volume, using the thickness of one water layer of 0.28 nm.
4. The local minima on the continuous density map define the centers of each LDA. A threshold of 33 nm^3^ (average water density, considering only the heavy atoms) was used to defines the border of the LDAs (Fig. 4D).
5. When neighboring LDAs were connected, we used a watershed algorithm to split them (Fig. 4E).
6. To sample the size, we started from the LDA’s center and for each sampled direction symmetrically extended outwards until one of the two ends intersected the LDA’s border, and used that as the maximum length in that direction. For each LDA, we sampled the lengths at 18 unique angles (equally spaced over 180° (Fig. 4F). All the pairs formed by perpendicular directions were considered as a width-and-height pair, and were used to build the final width-height distribution of LDAs (Fig. 5).

### GQD in Lipid Vesicle

#### GQD encapsulation

GQDs (2 nm to 8 nm in diameter) were encapsulated in lipid bilayer vesicles using a double emulsion generating microfluidic device as demonstrated previously.^59^ Briefly, the water/oil/water double emulsions were made by flow focusing with the inner solution containing the GQDs. The outer solution consisted of 20 mM K-HEPES (pH 7.4), 80 mM KCl, 10 mM MgCl_2_, 10% PVA (M.W. 13000 : 23000), 2% glycerol, 1% Poloxamer-188 and 250 mM glucose. The GQD sample was diluted in a buffer with final concentrations of 20 mM K-HEPES (pH 7.4), 80 mM KCl, 1 mM MgCl_2_, 2% PVA, 2% glycerol, 1% Poloxamer-188 and 180 mM sucrose to match the osmolarities of the solutions on both sides of the vesicle membrane. No significant difference in fluorescence intensity was observed for GQD samples diluted in water when compared to those diluted in the above-mentioned buffer. Lipids (POPC and 18:1 Liss Rhod PE) and cholesterol were purchased from Avanti Polar Lipids Inc. (Alabaster, AL). Liss Rhod PE was added at 0.01% by moles to impart fluorescence to the vesicle membrane for visualization. The formed double emulsion templated vesicles were collected in an imaging chamber made from sticking a glass coverslip on two double-sided tapes fixed onto a glass slide at a certain spacing. The demixing of middle organic phase took place within few minutes of their formation. A 64:36 percentage solution by volume of hexane and chloroform was used for dissolving the lipids as the middle phase.

#### Imaging

An Olympus IX-81 spinning disk confocal microscope was used to image the vesicles encapsulated with GQD samples. Fluorescence images using a GFP filter (ex-486 nm/em-515 nm) were captured at 500 ms exposure with a 60X oil objective (PlanApo, NA:1.42) at time interval of five minutes for 2 h. For vesicle visualization, a TRITC filter (ex-561 nm/em-610 nm) was used at an exposure of 100 ms.

#### Image analysis

All image analysis was carried out using ImageJ. Average fluorescence intensities were measured by drawing rectangular boxes within the interior of the vesicles at different places and taking their mean. Background intensity was calculated using the same approach but with the boxes drawn outside the vesicles in their immediate surroundings. Background subtracted intensities were then normalized with respect to the maximum intensity at time 0. A two-tailed student t-test was carried out with a significance level of 0.05 to compare the background subtracted intensities for the different lipid compositions between *t* = 0 and *t* =1 h.

## Abbreviations

COM, GQD, LDA, MD, NP, POPC

## Acknowledgement

This work was funded by the University of Michigan College of Engineerings Blue Sky Initiative. PE thanks the College of Engineering and Department of Mechanical Engineering at the University of Michigan for partial support.

## References

1. Mitragotri, S.; Burke, P. A.; Langer, R. Overcoming the challenges in administering biopharmaceuticals: Formulation and delivery strategies. Nature Reviews Drug Discovery 2014, 13, 655–672.

2. Jing, Y.; Zhu, Y.; Yang, X.; Shen, J.; Li, C. Ultrasound-triggered smart drug release from multifunctional core-shell capsules one-step fabricated by coaxial electrospray method. Langmuir 2011, 27, 1175–1180.

3. Chen, M. L.; He, Y. J.; Chen, X. W.; Wang, J. H. Quantum-dot-conjugated graphene as a probe for simultaneous cancer-targeted fluorescent imaging, tracking, and monitoring drug delivery. Bioconjugate Chemistry 2013, 24, 387–397.

4. Tian, Z.; Yao, X.; Ma, K.; Niu, X.; Grothe, J.; Xu, Q.; Liu, L.; Kaskel, S.; Zhu, Y. MetalOrganic Framework/Graphene Quantum Dot Nanoparticles Used for Synergistic Chemo- and Photothermal Therapy. ACS Omega 2017, 2, 1249–1258.

5. Liu, Z.; Robinson, J. T.; Sun, X.; Dai, H. PEGylated Nano-Graphene Oxide for Delivery of Water Insoluble Cancer Drugs-Supplementary Information. Journal of the American Chemical Society 2008, 130, 10876–10877.

6. Liu, Q.; Guo, B.; Rao, Z.; Zhang, B.; Gong, J. R. Strong two-photon-induced fluorescence from photostable, biocompatible nitrogen-doped graphene quantum dots for cellular and deep-tissue imaging. Nano Letters 2013, 13, 2436–2441.

7. Shi, W.; Fan, H.; Ai, S.; Zhu, L. Preparation of fluorescent graphene quantum dots from humic acid for bioimaging application. New Journal of Chemistry 2015, 39, 7054–7059.

8. Al-Nahain, A.; Lee, J. E.; In, I.; Lee, H.; Lee, K. D.; Jeong, J. H.; Park, S. Y. Target delivery and cell imaging using hyaluronic acid-functionalized graphene quantum dots. Molecular Pharmaceutics 2013, 10, 3736–3744.

9. Liu, J. J.; Zhang, X. L.; Cong, Z. X.; Chen, Z. T.; Yang, H. H.; Chen, G. N. Glutathione-functionalized graphene quantum dots as selective fluorescent probes for phosphate-containing metabolites. Nanoscale 2013, 5, 1810–1815.

10. Zhao, H.; Chang, Y.; Liu, M.; Gao, S.; Yu, H.; Quan, X. A universal immunosensing strategy based on regulation of the interaction between graphene and graphene quantum dots. Chemical Communications 2013, 49, 234–236.

11. Pridgen, E. M.; Alexis, F.; Farokhzad, O. C. Polymeric nanoparticle drug delivery technologies for oral delivery applications. Expert Opinion on Drug Delivery 2015, 12, 1459–1473.

12. Gelperina, S.; Kisich, K.; Iseman, M. D.; Heifets, L. The potential advantages of nanoparticle drug delivery systems in chemotherapy of tuberculosis. American Journal of Respiratory and Critical Care Medicine 2005, 172, 1487–1490.

13. Smith, J. D.; Morton, L. D.; Ulery, B. D. Nanoparticles as synthetic vaccines. Current Opinion in Biotechnology 2015, 34, 217–224.

14. Dames, P.; Gleich, B.; Flemmer, A.; Hajek, K.; Seidl, N.; Wiekhorst, F.; Eberbeck, D.; Bittmann, I.; Bergemann, C.; Weyh, T. et al. Targeted delivery of magnetic aerosol droplets to the lung. Nature Nanotechnology 2007, 2, 495–499.

15. Chattopadhyay, S. Aerosol generation using nanometer liposome suspensions for pulmonary drug delivery applications. Journal of Liposome Research 2013, 23, 255–267.

16. Beck-Broichsitter, M.; Merkel, O. M.; Kissel, T. Controlled pulmonary drug and gene delivery using polymeric nanocarriers. Journal of Controlled Release 2012, 161, 214–224.

17. Wadia, J. S.; Stan, R. V.; Dowdy, S. F. Transducible TAT-HA fusogenic peptide enhances escape of TAT-fusion proteins after lipid raft macropinocytosis. Nature Medicine 2004, 10, 310–315.

18. Zuris, J. A.; Thompson, D. B.; Shu, Y.; Guilinger, J. P.; Bessen, J. L.; Hu, J. H.; Maeder, M. L.; Joung, J. K.; Chen, Z. Y.; Liu, D. R. Cationic lipid-mediated delivery of proteins enables efficient protein-based genome editing in vitro and in vivo. Nature Biotechnology 2015, 33, 73–80.

19. Rabideau, A. E.; Pentelute, B. L. Delivery of Non-Native Cargo into Mammalian Cells Using Anthrax Lethal Toxin. ACS Chemical Biology 2016, 11, 1490–1501.

20. Luo, D.; Mark, W S. Synthetic DNA delivery systems. Nature biotechnology 2000, 18, 33–37.

21. Vader, P.; Mol, E. A.; Pasterkamp, G.; Schiffelers, R. M. Extracellular vesicles for drug delivery. Advanced Drug Delivery Reviews 2016, 106, 148–156.

22. Torchilin, V. P. Multifunctional, stimuli-sensitive nanoparticulate systems for drug delivery. Nature Reviews Drug Discovery 2014, 13, 813–827.

23. Overton, E. On the general osmotic properties of the cell, their probable origin,and their signicance for physiology (translated from German). Vierteljahrsschr Naturforsch Ges Zurich 1899, 44, 88–135.

24. Meyer, H. Which characteristic of the anesthetic causes its narcotic effect? (translated from German). Archiv für experimentelle Pathologie und Pharmakologie 1899, 42, 109–118.

25. Nagle, J. F.; Mathai, J. C.; Zeidel, M. L.; Tristram-Nagle, S. Theory of Passive Permeability through Lipid Bilayers. The Journal of General Physiology 2008, 131, 77–85.

26. Song, B.; Yuan, H.; Jameson, C. J.; Murad, S. Role of surface ligands in nanoparticle permeation through a model membrane: A coarse-grained molecular dynamics simulations study. Molecular Physics 2012, 110, 2181–2195.

27. Zocher, F.; Van Der Spoel, D.; Pohl, P.; Hub, J. S. Local partition coefficients govern solute permeability of cholesterol-containing membranes. Biophysical Journal 2013, 105, 2760–2770.

28. Parisio, G.; Stocchero, M.; Ferrarini, A. Passive membrane permeability: Beyond the standard solubility-diffusion model. Journal of Chemical Theory and Computation 2013, 9, 5236–5246.

29. Russ, K. A.; Elvati, P.; Parsonage, T. L.; Dews, A.; Jarvis, J. A.; Ray, M.; Schneider, B.; Smith, P. J.; Williamson, P. T.; Violi, A. et al. C60fullerene localization and membrane interactions in RAW 264.7 immortalized mouse macrophages. Nanoscale 2016, 8, 4134–4144.

30. Xiang, T. X.; Anderson, B. D. Influence of chain ordering on the selectivity of dipalmitoylphosphatidylcholine bilayer membranes for permeant size and shape. Biophysical Journal 1998, 75, 2658–2671.

31. Mathai, J. C.; Tristram-Nagle, S.; Nagle, J. F.; Zeidel, M. L. Structural Determinants of Water Permeability through the Lipid Membrane. The Journal of General Physiology 2008, 131, 69–76.

32. Xiang, T. X.; Anderson, B. D. Permeability of acetic acid across gel and liquid-crystalline lipid bilayers conforms to free-surface-area theory. Biophysical Journal 1997, 72, 223–237.

33. Mitragotri, S.; Johnson, M. E.; Blankschtein, D.; Langer, R. An analysis of the size selectivity of solute partitioning, diffusion, and permeation across lipid bilayers. Biophysical Journal 1999, 77, 1268–1283.

34. Bemporad, D.; Essex, J. W.; Luttmann, C. Permeation of small molecules through a lipid bilayer: a computer simulation study. J. Phys. Chem. 2004, B, 4875–4884.

35. Vamparys, L.; Gautier, R.; Vanni, S.; Bennett, W. D.; Tieleman, D. P.; Antonny, B.; Etchebest, C.; Fuchs, P. F. Conical Lipids in Flat Bilayers Induce Packing Defects Similar to that Induced by Positive Curvature. Biophysical Journal 2013, 104, 585–593.

36. Vance, J. E.; Vance, D. E. Elsevier; Elsevier, 2008.

37. Sun, H.; Gao, N.; Dong, K.; Ren, J.; Qu, X. Graphene quantum dots-band-aids used for wound disinfection. ACS Nano 2014, 8, 6202–6210.

38. Suzuki, N.; Wang, Y.; Elvati, P.; Qu, Z.-B.; Kim, K.; Jiang, S.; Baumeister, E.; Lee, J.; Yeom, B.; Bahng, J. H. et al. Chiral Graphene Quantum Dots. ACS nano 2016, 10, 1744–55.

39. Barry, P. H.; Diamond, J. M. Effects of unstirred layers on membrane phenomena. Physiological Reviews 1984, 64, 763–872.

40. Tu, Y.; Lv, M.; Xiu, P.; Huynh, T.; Zhang, M.; Castelli, M.; Liu, Z.; Huang, Q.; Fan, C.; Fang, H. et al. Destructive extraction of phospholipids from Escherichia coli membranes by graphene nanosheets. Nature Nanotechnology 2013, 8, 594–601.

41. Chang, R.; Violi, A. Insights into the effect of combustion-generated carbon nanoparticles on biological membranes: A computer simulation study. Journal of Physical Chemistry B 2006, 110, 5073–5083.

42. Duan, G.; Zhang, Y.; Luan, B.; Weber, J. K.; Zhou, R. W.; Yang, Z.; Zhao, L.; Xu, J.; Luo, J.; Zhou, R. Graphene-Induced Pore Formation on Cell Membranes. Scientific Reports 2017, 7, 42767.

43. Zucker, I.; Werber, J. R.; Fishman, Z. S.; Hashmi, S. M.; Gabinet, U. R.; Lu, X.; Osuji, C. O.; Pfefferle, L. D.; Elimelech, M. Loss of Phospholipid Membrane Integrity Induced by Two-Dimensional Nanomaterials. Environmental Science and Technology Letters 2017, 4, 404–409.

44. Leroueil, P. R.; Berry, S. A.; Duthie, K.; Han, G.; Rotello, V. M.; McNerny, D. Q.; Baker, J. R.; Orr, B. G.; Holl, M. M. B. Wide varieties of cationic nanoparticles induce defects in supported lipid bilayers. Nano Letters 2008, 8, 420–424.

45. Ong, S.; Liu, H.; Pidgeon, C. Immobilizedartificial-membrane chromatography: measurements of membrane partition coefficient and predicting drug membrane permeability. Journal of Chromatography A, 1996, 728, 113–128.

46. Wohnsland, F.; Faller, B. High-throughput permeability pH profile and high-throughput alkane/water log P with artificial membranes. Journal of Medicinal Chemistry 2001, 44, 923–930.

47. Garrido, N. M.; Queimada, A. J.; Jorge, M.; Macedo, E. A.; Economou, I. G. 1-Octanol / Water Partition Coefficients of n -Alkanes from Molecular Simulations of Absolute Solvation Free. Journal of Chemical Theory and Computation 2009, 5, 2436–2446.

48. Moriguchi, I.; Shuichi, H.; Liu, Q.; Nakagome, I.; Matsushita, Y. Simple method of calculating octanol/water partition coefficient. Chemical and Pharmaceutical Bulletin 1992, 40, 127–130.

49. Meylan, WM; Howard, P. Atom/fragment contribution method for estimating octanol-water partition coefficients. -PubMed - NCBI. Journal of Pharmaceutical Sciences 1995, 84, 83–92.

50. Walt, v. d. S.; Sch, J. L.; Nunez-iglesias, J.; Francois Boulogne,; Warner, J. D.; Yager, N.; Gouillart, E.; Yu, T. scikitimage: image processing in Python. PeerJ 2014, 2e, 1–18.

51. Peliti, L.; Leibler, S. Effects of thermal fluctuations on systems with small surface tension. Physical Review Letters 1985, 54, 1690–1693.

52. Brown, M. F.; Ribeiro, A. A.; Williams, G. D. New view of lipid bilayer dynamics from 2H and 13C NMR relaxation time measurements. Proceedings of the National Academy of Sciences 1983, 80, 4325–4329.

53. Hayduk, W.; Buckley, W. D. Effect of molecular size and shape on diffusivity in dilute liquid solutions. Chemical Engineering Science 1972, 27, 1997–2003.

54. Cui, H.; Lyman, E.; Voth, G. A. Mechanism of membrane curvature sensing by amphipathic helix containing proteins. Biophysical Journal 2011, 100, 1271–1279.

55. Almeida, P. F.; Thompson, T. E. Lateral Diffusion in the Liquid Phases of Dimyristoylphosphatidylcholine/Cholesterol Lipid Bilayers: A Free Volume Analysis. Biochemistry 1992, 31, 6739–6747.

56. Hung, W. C.; Lee, M. T.; Chen, F. Y.; Huang, H. W. The condensing effect of cholesterol in lipid bilayers. Biophysical Journal 2007, 92, 3960–3967.

57. Pan, J.; Mills, T. T.; Tristram-Nagle, S.; Nagle, J. F. Cholesterol perturbs lipid bilayers nonuniversally. Physical Review Letters 2008, 100, 1–4.

58. Issack, B. B.; Peslherbe, G. H. Effects of Cholesterol on the Thermodynamics and Kinetics of Passive Transport of Water through Lipid Membranes. Journal of Physical Chemistry B 2015, 119, 9391–9400.

59. Majumder, S.; Garamella, J.; Wang, Y. L.; Denies, M.; Noireaux, V.; Liu, A. P. Cell-sized mechanosensitive and biosensing compartment programmed with DNA. Chemical Communications 2017, 53, 7349–7352.

60. Phillips, J. C.; Braun, R.; Wang, W.; Gumbart, J.; Tajkhorshid, E.; Villa, E.; Chipot, C.; Skeel, R. D.; Kalé, L.; Schulten, K. Scalable molecular dynamics with NAMD. Journal of Computational Chemistry 2005, 26, 1781–1802.

61. Klauda, J. B.; Venable, R. M.; Freites, J. A.; Connor, J. W. O.; Tobias, D. J.; Mondragon-ramirez, C.; Vorobyov, I.; Mackerell, A. D.; Pastor, R. W. Update of the CHARMM all-atom additive force field for lipids: validation on six lipid types. physical chemistry B 2010, 2, 7830–7843.

62. Vanommeslaeghe, K.; Hatcher, E.; Acharya, C.; Kundu, S.; Zhong, S.; Shim, J.; Darian, E.; Guvench, O.; Lopes, P.; Vorobyov, I. et al. CHARMM general force field: A force field for drug-like molecules compatible with the CHARMM all-atom additive biological force fields. Journal of Computational Chemistry 2010, 31, 671–690.

63. MacKerell, A. D.; Bashford, D.; Bellott, M.; Dunbrack, R. L.; Evanseck, J. D.; Field, M. J.; Fischer, S.; Gao, J.; Guo, H.; Ha, S. et al. All-Atom Empirical Potential for Molecular Modeling and Dynamics Studies of Proteins. The Journal of Physical Chemistry B 1998, 102, 3586–3616.

64. Vanommeslaeghe, K.; MacKerell, A. D. Automation of the CHARMM general force field (CGenFF) I: Bond perception and atom typing. Journal of Chemical Information and Modeling 2012, 52, 3144–3154.

65. Jo, S.; Kim, T.; Im, W. Automated builder and database of protein/membrane complexes for molecular dynamics simulations. PLoS ONE 2007, 2.

66. Ryckaert, J. P.; Ciccotti, G.; Berendsen, H. J. Numerical integration of the cartesian equations of motion of a system with constraints: molecular dynamics of n-alkanes. Journal of Computational Physics 1977, 23, 327–341.

67. Darden, T.; York, D.; Pedersen, L. Particle mesh Ewald: An N·log(N) method for Ewald sums in large systems. The Journal of Chemical Physics 1993, 98, 10089–10092.

68. Loncharich, R. J.; Brooks, B. R.; Pastor, R. W. Langevin dynamics of peptides: The frictional dependence of isomerization rates of NacetylalanylNmethylamide. Biopolymers 1992, 32, 523–535.

69. Martyna Tobias, K. Constant pressure molecular dynamics algorithms — Issue 5-Journal of Chemical Physics. Journal of Chemical Physics 1994, 6323.

70. Feller, S. E. Constant pressure molecular dynamics simulation-the langevin piston method. J. Comput. Chem. 1995, 103, 4613–4621.

71. Humphrey, W.; Dalke, A.; Schulten, K. VMD - Visual Molecular Dynamics. Journal of Molecular Graphics 1996, 14, 33–38.

72. Michaud-Agrawal, N.; Denning, E. J.; Woolf, T. B.; Beckstein, O. MDAnalysis: A Toolkit for the Analysis of Molecular Dynamics Simulations. Journal of computational chemistryomputational chemistry 2011, 32, 2319–2327.

73. Gowers, R. J.; Linke, M.; Barnoud, J.; Reddy, T. J. E.; Melo, M. N.; Seyler, S. L.; Domanski, J.; Dotson, D. L.; Buchoux, S.; Kenney, I. M. et al. MDAnalysis: A Python Package for the Rapid Analysis of Molecular Dynamics Simulations. Proceedings of the 15th Python in Science Conference 2016, Scipy, 98–105.

