## Supporting Information for "Predicting the time of entry of nanoparticles in cellular membranes"

### Supporting Information
### Predicting the time of entry of nanoparticles in cellular membranes

Changjiang Liu[a], Paolo Elvati[b], Sagardip Majumder[b], Yichun Wang[c], Allen Liu[a,b], Angela Violi[a,b,c,d]

[a]*Department of Biophysics, University of Michigan, Ann Arbor, MI 48109-2125, USA*
[b]*Department of Mechanical Engineering, University of Michigan, Ann Arbor, MI 48109-2125, USA*
[c]*Departments of Chemical Engineering, University of Michigan, Ann Arbor, MI 48109-2125, USA*
[d]*Biomedical Engineering, Macromolecular Science and Engineering, Biophysics Program, University of Michigan, Ann Arbor, MI, United States*

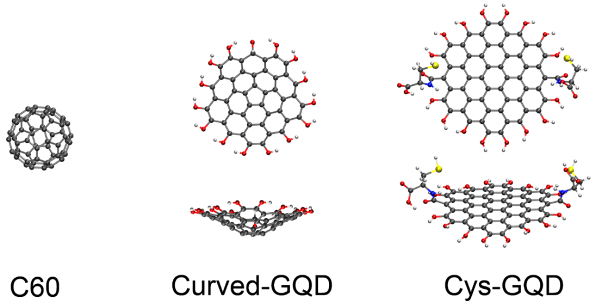


Figure S1: C60 as shown in the picture. The curved-GQD had a five-member ring at the center and hydroxyl groups on the surface. The cys-GQD had two L-cysteines had hydroxyl groups on the surface except the 2, 11 which had each a L-cysteine connected by covalent bond.

*Corresponding author:

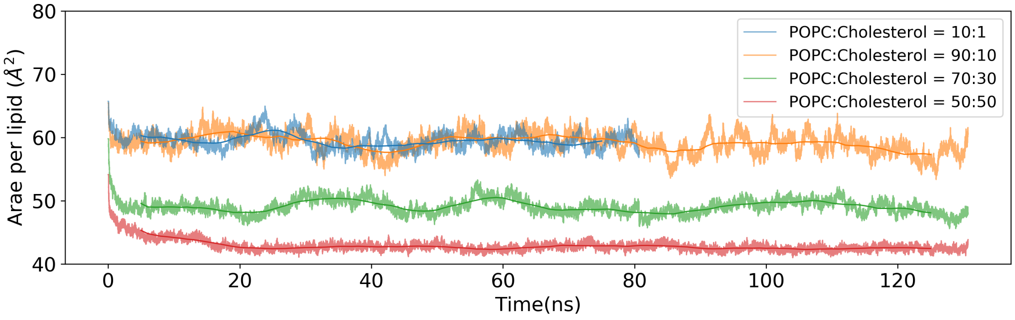


Figure S2: The area per lipid over time for the four types of membrane presented in this work. Light lines represent the area per lipids over time for each frame. Dark lines represent the running average of the light line over a 10 ns window. The relaxation of the POPC:cholesterol = 10:1 membrane was done by 70 ns simulations. The relaxation of the other three membranes (POPC:cholesterol = 90:10, 70:30 and 50:50) were done by 130 ns simulations. The membranes were assumed at equilibrium at the tailing 50 ns as time average of the area per lipid in a NPsT simulations was varying less than 10%.

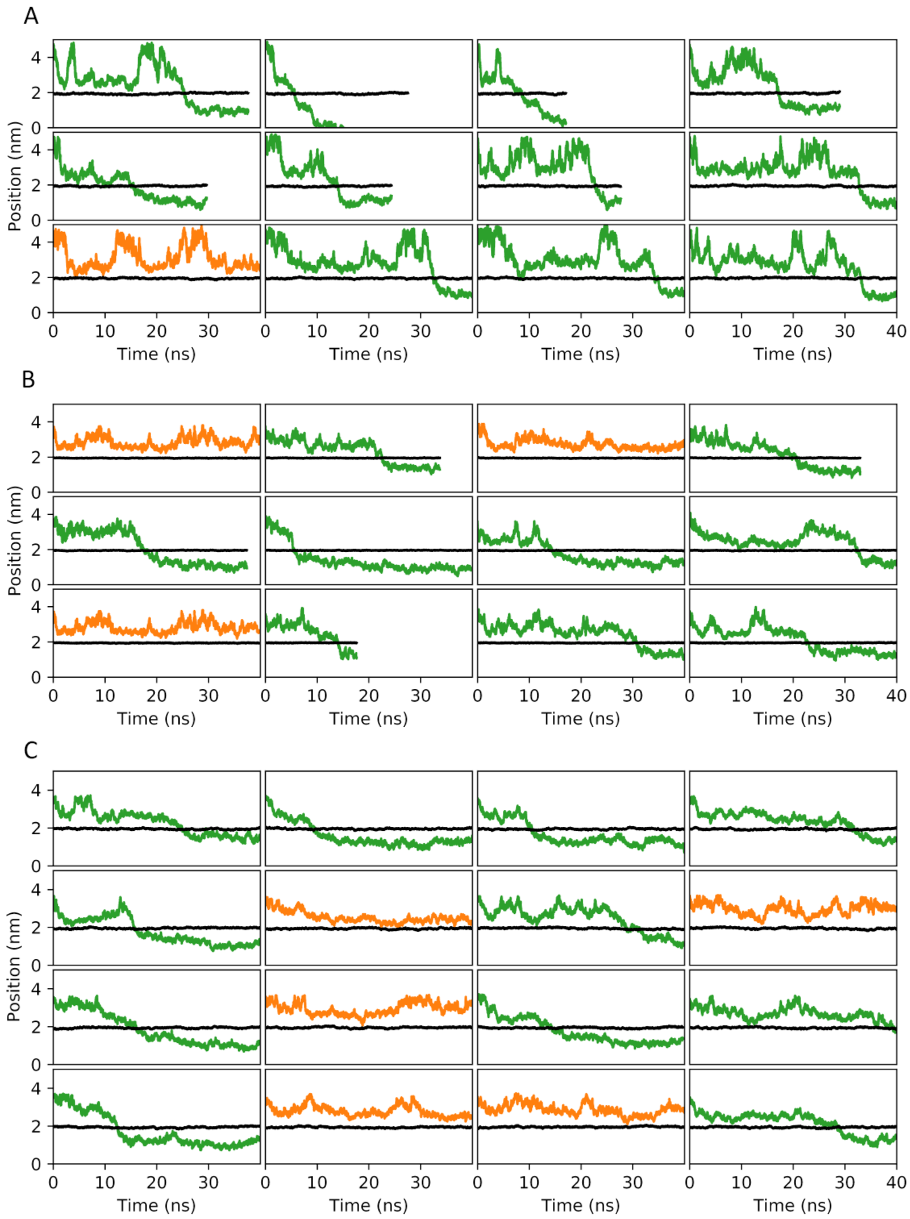


Figure S3: (A) 12 simulations of C60 with POPC/cholesterol membrane. (B) 12 simulations of curved-GQD with POPC/cholesterol membrane. (C) 16 simulations of cys-GQD with POPC/cholesterol membrane. Green (that entered) and orange (that did not enter) trajectories showed where the C60s were regarding the membrane along the z-axis. The black line showed the surface of the membrane for comparison.
